# Immunization by exposure to live virus (SIV_mne_/HIV-2_287_) during antiretroviral drug prophylaxis reduces risk of subsequent viral challenge

**DOI:** 10.1101/2020.09.30.320820

**Authors:** LM Frenkel, L Kuller, IA Beck, C-C Tsai, JP Joy, TM Mulvania, DC Montefiori, DM Anderson

## Abstract

**Rationale/Study Design:** A major challenge in the development of HIV vaccines is finding immunogens that elicit protection against a broad range of viral strains. Immunity to a narrow range of viral strains may protect infants of HIV-infected women or partners discordant for HIV. We hypothesized that immunization to the relevant viral variants could be achieved by exposure to infectious virus during prophylaxis with antiretroviral drugs. To explore this approach in an animal model, macaques were exposed to live virus (SIV_mne_ or HIV-2_287_) during prophylaxis with parenteral tenofovir. The humoral and cellular immune responses were quantified. Subsequently, experimental animals were challenged with homologous virus to evaluate protection from infection, and if infection occurred, the course of disease was compared to control animals. Experimental animals uninfected with SIV_mne_ were challenged with heterologous HIV-2_287_ to assess resistance to retroviral infection.

**Methodology/Principal Findings:** Juvenile *Macaca nemestrina* (N=8) were given ten weekly intravaginal exposures with either moderately (SIV_mne_) or highly (HIV-2_287_) pathogenic virus during tenofovir prophylaxis. Tenofovir protected all 8 experimental animals from infection, while all untreated control animals became infected. Specific non-neutralizing antibodies were elicited in blood and vaginal secretions of experimental animals, but no ELISPOT responses were detected. Six weeks following the cessation of tenofovir, intravaginal challenge with homologous virus infected 2/4 (50%) of the SIV_mne_-immunized animals and 4/4 (100%) of the HIV-2_287_-immunized animals. The two SIV_mne_-infected and 3 (75%) HIV-2_287_-infected had attenuated disease, suggesting partial protection.

**Conclusions/Significance:** Repeated exposure to SIV_mne_ or HIV-2_287_ during antiretroviral prophylaxis blocked infection induced binding antibodies in the blood and mucosa, but not neutralizing antibodies or specific cellular immune responses. Studies to determine whether antibodies are similarly induced in breastfeeding infants and sexual partners discordant for HIV infection and receiving pre-exposure antiretroviral prophylaxis are warranted, including whether these antibodies appear to confer partial or complete protection from infection.

## Introduction

The concentration of human immunodeficiency virus type-1 (**HIV**) virions in an infected individual’s blood and genital secretions correlates with viral transmission to his or her sexual partner [1]. Likewise, HIV levels in blood and breastmilk correlate with transmission from an infected woman to her infant [2, 3], as do multiple other factors (e.g., genital infections [4, 5], progestins (reviewed [6]), and providing complementary foods during breastfeeding [7]). Certain human genetic variants and polymorphisms can reduce the likelihood of HIV acquisition, including deletions in the gene that encodes the HIV co-receptor CCR5 [8] and other less clearly defined factors (e.g., mucosal and breast milk non-specific antiviral proteins, enzymes, and cells) [9, 10].

Given that HIV causes a persistent infection that without treatment has high mortality, considerable effort has been devoted to the development of a vaccine that can prevent infection (reviewed [11]). Initial efforts to generate a protective HIV vaccine were largely focused on eliciting protective immunity via broadly neutralizing antibodies (reviewed [12]). This approach was pursued due to observations that sera of chronically HIV-infected individuals neutralized significant numbers of heterologous virus isolates [13], and the transfer of sera containing neutralizing HIV/SIV antibodies and neutralizing monoclonal antibodies have protected experimental animals from mucosal challenge [14-16]. However, the breadth of vaccine-elicited neutralizing antibodies has generally been narrow (reviewed [17]). Also problematic is that in most individuals, the induction of broadly neutralizing antibodies requires a series of mutations in V-beta chains that occurs over months to years [18].

Modest protection from HIV infection was achieved in one vaccine trial (RV144) in humans with two doses of ALVAC-HIV, a canarypox vector expressing HIV-1 Env, Gag, Protease, and a transmembrane region of gp41, followed by two boosts with ALVAC-HIV plus AIDSVAX clades B/E gp120 protein [19]. Protection from infection was associated with the generation of IgG binding antibodies to the V2 Env loop [20, 21]. As neutralizing antibodies and cellular responses were rarely detected in this trial [22], these binding antibodies likely conferred protection from HIV acquisition. Due to the low overall protective effect estimated at 31% [19], another trial of this vaccine strategy was initiated but recently halted due to futility.

Many individuals at-risk of HIV infection, particularly breastfeeding infants of infected mothers and sexual partners discordant for HIV infection, are repeatedly exposed to the same viral swarm from mother’s breast milk or his/her infected partner’s genital fluids. These individuals could potentially benefit from a “narrow” immune response to the HIV swarm during repeat viral exposures. We hypothesized that a specific immune response could be induced to the relevant virus and protect the susceptible individual from HIV infection. In support of this are the observations that: (1) Seronegative sexual partners of HIV-infected individuals with undetectable plasma viral loads develop HIV-specific CD8+ T cell responses, suggesting that low but persistent, exposure to HIV may be sufficient to elicit virus-specific immune responses [23, 24] and (2) The combination of an Env vaccine and pre-exposure prophylaxis (tenofovir microbicide gel) reduced the risk of HIV infection more than either intervention alone [25]. Taken together, repeated viral exposures during pre-exposure prophylaxis with antiretrovirals may induce protective immune responses to a specific viral quasispecies.

Pre-exposure prophylaxis (PrEP) with antiretrovirals is recommended for individuals at risk of HIV infection in the US and globally based on several clinical trials (CDC & WHO, 2015) [26-29]. Individuals on PrEP can develop mucosal HIV-1-specific IgA immune responses [30], which may protect them from HIV acquisition.

The current study explores in a controlled trial of macaques whether multiple exposures to infectious virus during antiretroviral prophylaxis induces specific immunity to the relevant viruses by repeated exposures during antiretroviral prophylaxis. Each animal’s binding and neutralizing antibodies and cellular immune responses were measured. Subsequently, the macaques were challenged with homologous virus. If uninfected after three homologous viral challenges, then a heterologous viral challenge was administered to determine if the animal was susceptible to retroviral infection. Infected animals were monitored to evaluate whether the disease course was attenuated compared to controls.

## Methods

### Study animals and design

The animals used for study were pre-pubertal female *Macaca nemestrina*, clinically healthy by physical examination, serum chemistries, and complete blood counts, including negative serology and polymerase chain reaction assay (**PCR**) of peripheral blood mononuclear cells (**PBMC**) for simian retrovirus (**SRV**) infection. The macaques were housed singly in stainless steel cages in the Animal Biosafety Level 2-Plus facilities at the accredited Washington National Primate Research Center (**WaNPRC**). Animal care and husbandry met or exceeded the standards described in “The Guide for the Care and Use of Laboratory Animals” published by the Institute of Laboratory Animal Resources Commission on Life Sciences, National Research Council. All procedures were approved by the University of Washington’s Institutional Animal Care and Use Committee.

The study design (**Figure 1**) included administration of tenofovir, ({[(2*R*)-1-(6-amino-9*H*-purin-9-yl)propan-2-yl]oxy}methyl)phosphonic acid, 20 mg/kg by subcutaneous injection to experimental macaques once daily. The control animals were injected with normal saline as a placebo. The experimental and control macaques were inoculated intra-vaginally once a week for 10 weeks with 10^4^ tissue culture infectious dose for 50% of wells (**TCID**_**50**_) of virus, which was estimated to represent ten 100% animal infectious doses grown in autologous PBMC. The virus was introduced through a soft plastic 2.5 Fr nasal/gastric feeding tube into the vaginal vault followed by a flush of 1 mL sterile PBS.

**Figure 1.**
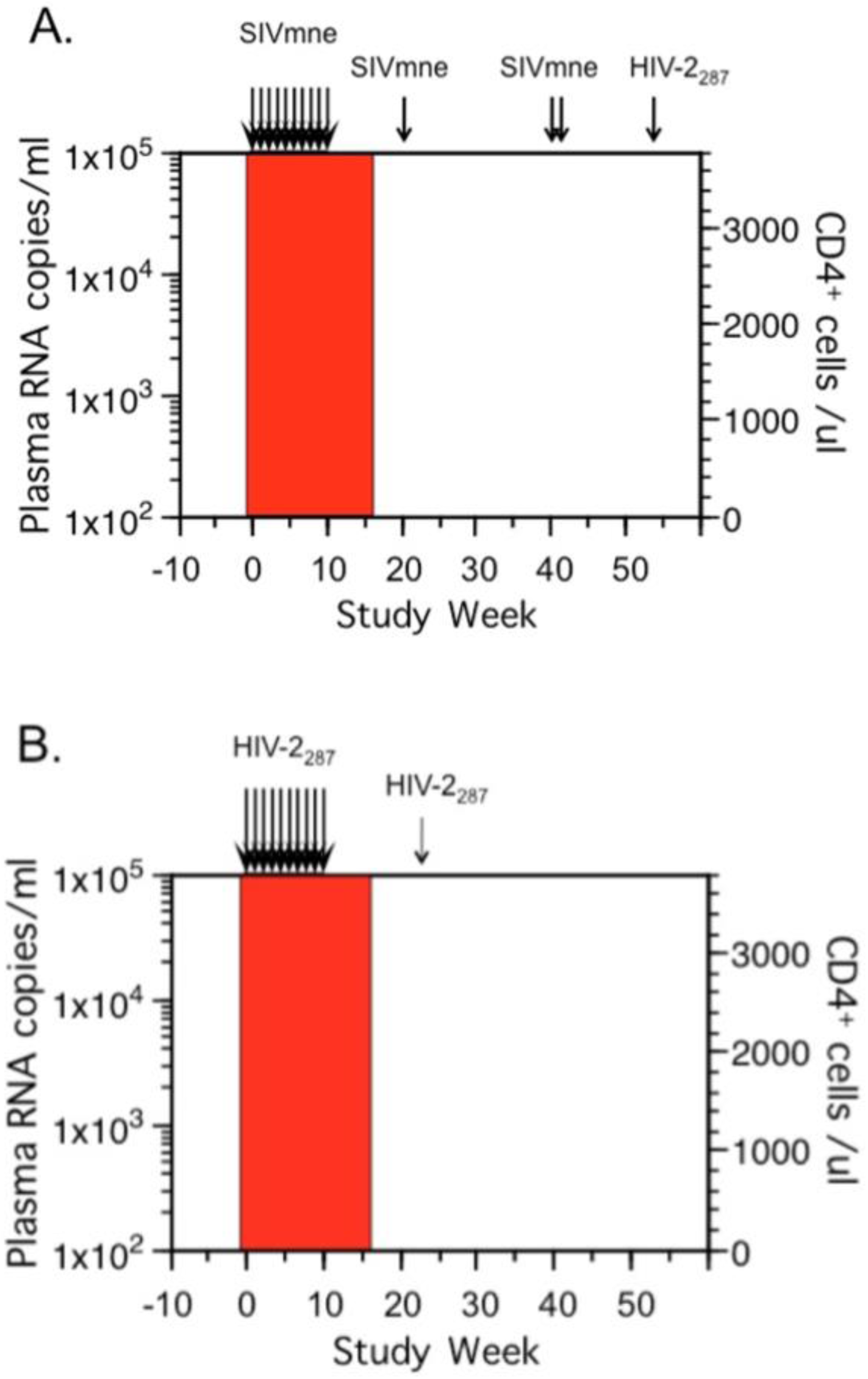
Study design. **A**. Panel A shows the time schema for immunization of experimental group (n=4) of female juvenile *Macaca nemestrina* given ten weekly intravaginal exposures to SIV_mne_ (multiple arrows on left) during tenofovir (red block). Controls did not receive tenofovir and were similarly inoculated (n=2 animals) or given only two inoculations 24 hours apart (n=2 animals). The virus used for each animal was grown in that animal’s own cells to avoid viral clearance based on allo-antigens incorporated into viral envelope. Macaques were intravaginally challenged initially (at Study Week 20) with 10^4^ TCID_50_ of SIV_mne_ grown in homologous cells (estimated 10 x the 100% animal infectious dose). Uninfected animals were subsequently challenged (at Study Week 41) with SIV_mne_ (two doses of 10^4.5^ TCID_50_ grown in allogenic animals and given 24 hours apart). To determine if uninfected animals were susceptible to retrovirus infection, they were inoculated with HIV-2_287_ (at Study Week 54). Blood and vaginal secretions were collected weekly over ten weeks, then every 2-6 weeks, and inguinal lymph nodes collected at Study Week 19. Specimens were assessed for viral RNA, DNA, infectivity, binding and neutralizing antibodies and cellular responses to Gag. **B**. Panel B shows the time schema for immunization and challenges of HIV-2_287_ experimental group (n=4). The study designed mirrored that in Panel A, except that the control animals (n=4) were only given two intravaginal exposures, and the experimental animals were only given one intravaginal challenge (at Study Week 20).

### Generation of viral stocks in each animal’s PBMC

Virus stocks were generated for each animal using autologous PBMCs to avoid viral clearance on the basis of macaque allo-antigens incorporated into the viral envelope [31, 32].

Phytohemoglutinin-A (**PHA**) stimulated CD8-depleted PBMCs from each macaque were infected with SIV_mne_ working stock B2455-1b or HIV-2_287_ working-stock-8 [33]. The cultures were monitored for cytopathic effects, specifically syncitia formation, and p27 antigen (**Ag**) production. At the peak of syncitia and p27Ag expression, uninfected CD8-depleted PBMCs from each macaque were independently added to the virus culture to allow a second phase of viral replication. The supernatant was subsequently filtered with a Centricon Plus-80 with Biomax-PB membrane (NMWL100,000) (Millipore, Bedford, MA) and stored in liquid nitrogen to be used as viral stock for each macaque made from her own PBMCs. To estimate the infectious titer of the stock viruses, a ten-fold dilution series using one stock aliquot each were carried out using 10^6^ macaque CD8-depleted PBMCs in quadruplicate across a 48-well plate. After maintenance of the cultures for 10 days, the viral antigen in the supernatant was measured and the TCID_50_/mL of the stock calculated.

### SIV/HIV-2 cultures of PBMCs with and without depletion of CD8^+^ T cells

The human CD4 T/B hybrid cell line, CEMx174, was used for SIV isolation [34], and CD8-depleted human PBMCs for HIV-2 isolation [35]. Two successive increases in antigen supernatant levels were used to define a positive culture. The number of HIV-2 positive cells were determined and reported as infectious units per million cells (**IUPM**).

### Quantitative PCR for SIV and HIV-2 plasma RNA

Plasma viral SIV RNA levels were determined using quantitative PCR (Bayer, Emeryville, CA) with a lower limit of detection of 1,000 copies/mL of plasma [36]. HIV-2 RNA plasma levels were quantified as described [35]. The lower limit of quantification of the HIV-2 assay was 1580 copies/mL.

### SIV/HIV-2 DNA PCR

Total DNA was extracted from isolated PBMC or tissues after grinding with mortar and pestle using the IsoQuick Nucleic Acid Extraction Kit (Orca Research, Bothell, WA). Nested PCR was performed on 2 ug of DNA (equivalent to ∼300,000 cells) per reaction utilizing primers specific for the LTR-gag region of SIV_mne_ and HIV-2_287_. The primers used were G1/G2B and S1/G4 (**G1**: TCTCTCCTAGTCGCCGCCTGGT; **G2B**: TTCATTTGCTGCCCATACTACA; **S1**: CGATAATAAGAAGACCCTGGTCTG and **G4**: TTCTTCCCTGACAAGACGGAG). The first-round cycling conditions were: 40 cycles of 94°C for 30 sec, 57°C for 30 sec and 72°C for 1 min, followed by an extension step of 72°C for 7 minutes. Two uL of first round product was used in the second round, the second-round cycling conditions were: 35 cycles of 94°C for 30 sec, 57°C for 30 sec and 72°C for 1 min, followed by an extension step of 72°C for 7 minutes. The 320-bp DNA fragment amplified was visualized by electrophoresis on 2% agarose gels and ethidium bromide staining. The limit of detection was 3-10 copies/10^6^ cells.

### Differentiation between infection by SIV_mne_ or HIV-2_287_

The viral nucleic acids detected by PCR of the LTR-*gag* region were digested with the restriction endonuclease ScaI (New England BioLabs, Beverly, MA), which cuts within the amplicon of HIV-2_287_ but not SIV_mne_. Five uL of each PCR product were digested with ScaI for 6 hours at 37°C. The digestion products, two DNA fragments of 240 and 80-bp for HIV-2 or one fragment of 320-bp for SIV, were separated on a 3% agarose gel and stained with ethidium bromide.

### Lymphocyte subset determinations in macaques

Absolute counts and ratios of CD4+ and CD8+ cells in the peripheral blood of virus-infected macaques were determined by standard fluorescent activated cell sorting (**FACS**) procedures using antibodies previously shown to react with macaque cells [34, 37].

### IFN-γ ELISpot to Gag

Functional CD8 T-cell activity was measured and quantified using a cytokine ELISpot assay [38, 39]. MHC Class I antigen presenting cells were constructed by infection of autologous *H. papio* transformed B cells with recombinant vaccinia viruses expressing *gag* of SIV, and combined with PBMCs at an effector to target ratio (**E:T**) of 2:1. Following an 18 hour incubation at 37°C, the cells were transferred in duplicate 2-fold dilutions from 2×10^5^-2.5×10^4^ cells/mL into an ELISpot plate that was pre-coated with monoclonal antibodies directed against macaque IFN-γ, and the assay done following the manufacturer’s instructions (U-Cytech, Utrecht, the Netherlands). The IFN-γ captured on the plate was detected by labeling with biotinylated polyclonal anti-IFN-γ antibodies followed by ϕ-labeled anti-biotin antibodies, and a proprietary activator that allows “spot” formation. The spots were counted using an ELISpot plate reader (Zeiss, Thornwood, New York). The average number of spots per well was compared to the number of input T cells, and linear regression used to determine the strength of the relationship made by the four data points (R^2^ value). The slope of the line indicated the frequency of IFN-γ secreting cells. The “partial F test” was utilized to determine if the experimental line was different from the control line.

### Quantitative total IgG antibody ELISA

Total antibody isotypes were determined using published methods [40]. A standard curve consisting of known amounts of purified IgG (Sigma, St. Louis MO) was run in conjunction with each assay. IgG levels in the samples were quantitatively interpolated from the standard curve by applying the linear regression equation obtained from the standard curve to the optical density values of the experimental samples. The reproducibility of the quantitative isotype-specific ELISA assay was evaluated by comparing the linear portion of both the experimental and standard sample curves (n=38). The standard deviation (**SD**) ranged from 1.3 x 10^−6^ to 3.6 x 10^−8^ between the observed antibody concentrations (experimental curves) and the known antibody concentrations (standard curves). A coefficient of variance (**CV**) of 1% to 5% indicated that the assay performed consistently.

### Quantitative SIV_mne_ and HIV-2_287_ antibody

Determination of virus-specific antibody levels was performed as previously described [41]. Virus-specific IgG levels were derived from the standard curve (run concurrently) as described above for the total IgG ELISA. The lowest level of detection for the quantitative ELISA is 1 x 10^−5^ µg antibody/mL. Reproducibility of the ELISA was evaluated by measuring the mid-point optical density of the linear portion of the standard curve for each assay (n=20). The SD for the SIV_mne_-specific ELISA was 0.072 and 0.066 for the HIV-2_287_ ELISA. A CV of 8.2% indicated that the assays performed consistently.

### Neutralizing Antibodies to SIV_mne_ or HIV-2_287_

Neutralizing antibodies against SIV_mne_ or HIV-2_287_ were assessed in CEMx174 cells as described previously [42]. Briefly 50 µL of cell-free virus (5,000 TCID_50_) was added to multiple dilutions of test serum in 100 µL of growth medium in triplicate wells of 96-well microtiter plates and incubated for 1 hr at 37°C. Cells (7.5 x 10^4^) in 100 µL of growth medium were added and incubated until extensive syncytium formation and nearly complete cell-killing were evident microscopically in virus control wells. Cell densities were reduced and medium replaced after 3 days incubation when more than 3 days were required to reach the assay end-point. Viable cells were stained with Finter’s neutral red in poly-L-lysine coated plates. Percent protection from virus-induced cell-killing was determined by calculating the difference in absorption (A_540_) between test wells (cells + serum sample + virus) and virus control wells (cells + virus), dividing this result by the difference in absorption between cell control wells (cells only) and virus control wells and multiplying by 100. Neutralizing antibody titers are expressed as the reciprocal of the serum dilution that protected 50% of cells from virus-induced killing. This cut-off corresponds to an approximate 90% reduction in p24 antigen synthesis. Assay stocks of virus were produced in either H9 cells (SIV_mne_) or human PBMC (HIV-2_287_).

## Results

### Tenofovir prophylaxis protected animals from infection

Antiretroviral prophylaxis with tenofovir given immediately prior to or at 24 hours following each of ten weekly intravaginal exposures to HIV-2_287_ or SIV_mne_ protected all 8 macaques from infection. In comparison, all placebo-treated controls became infected (**Figure 2**). Weekly monitoring for viral DNA were negative for all the experimental macaques’ PBMC at study weeks 1-14 during tenofovir administration (multiple PBMC aliquots totaled 14-30 µg DNA/animal). After cessation of tenofovir, multiple PBMC specimens were negative for virus by co-culture and PCR (6-10µg DNA/animal collected over study weeks 15 to 20) as was inguinal lymphoid tissue collected at study week 19 (total of 20-30 µg DNA/animal, each PCR with 2 µg DNA).

**Figure 2.**
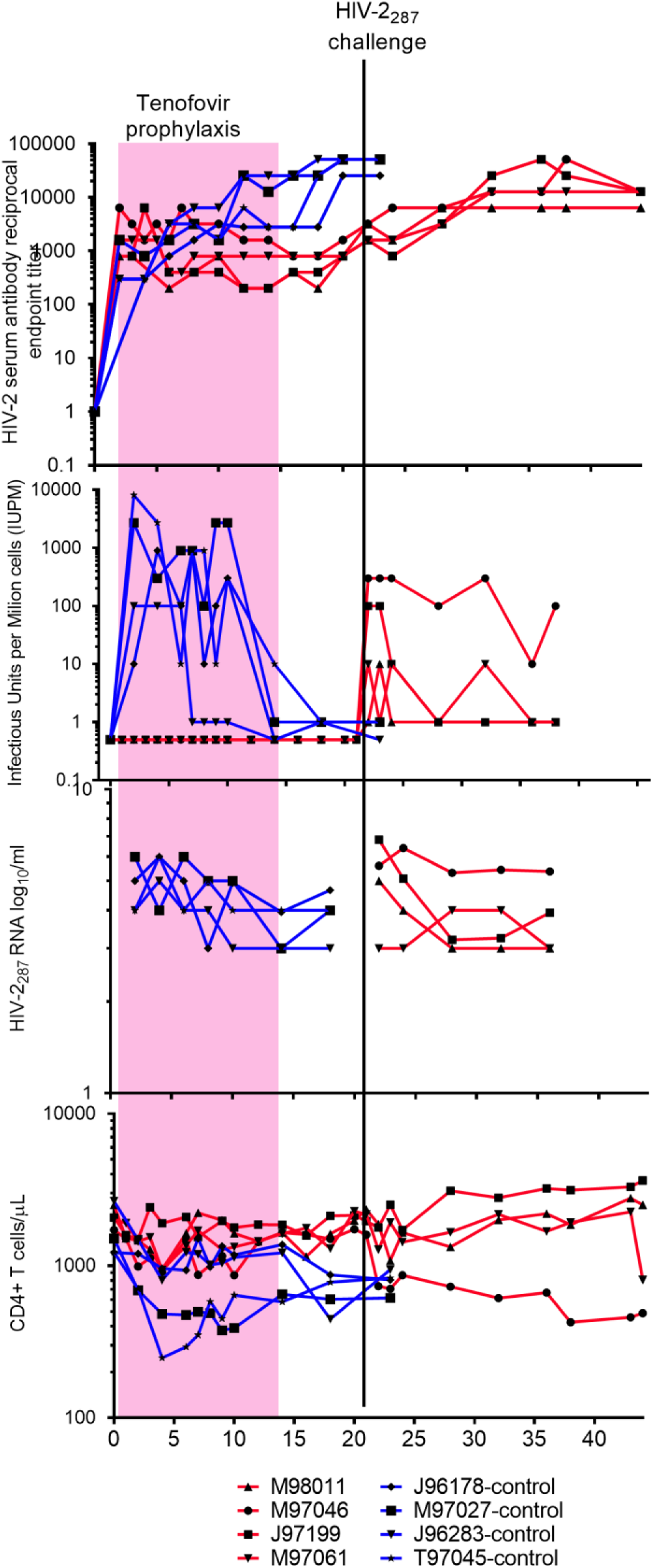
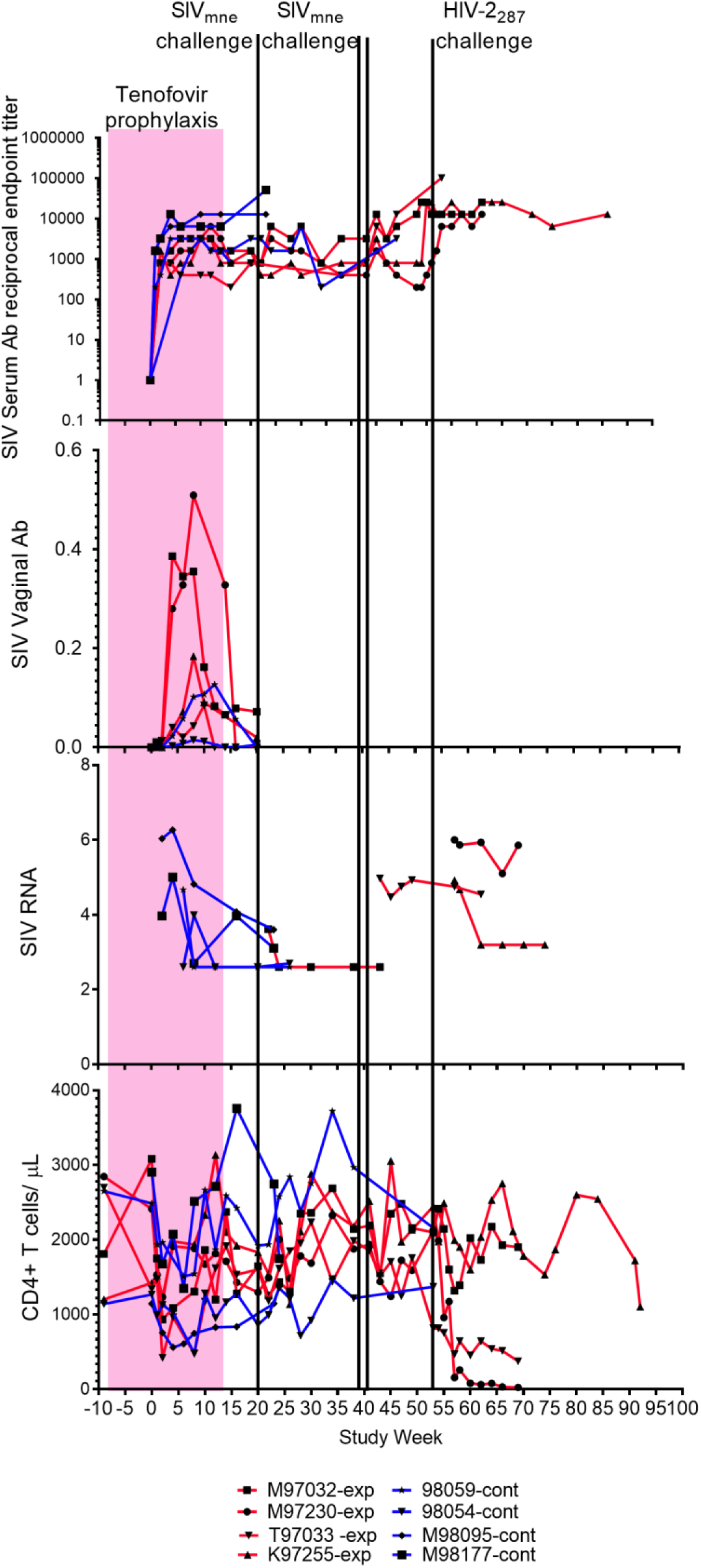
Macaques immunized to SIV/HIV-2 by exposure to virus during tenofovir prophylaxis resisted viral challenge and demonstrated protection from disease. **A**. Panels show SIV serum (top) and vaginal (second from top) antibody levels, viral load (third from top), and CD4 cell concentration (bottom) for control (blue lines) and experimental (red lines) macaques. Tenofovir (pink block) was given to experimental group from Study Week 0-14. Experimental animals had intravaginal challenges with SIV_Mne_ (green lines). One experimental animal, M97032, was infected by the first challenge of 10^4^ TCID_50_ of SIV_mne_ grown in homologous cells. A second animal, T97033, was infected by the second/third challenge with two doses of 10^4.5^ TCID_50_ SIV_mne_ grown in heterologous cells separated by 24 hours. Two control animals (M98095 and M98177) similarly received two intravaginal inoculations from this viral pool. The remaining two experimental animals, M97230 and K97255, were not infected by SIV_mne_ challenges; but were not resistant to retroviruses as infected by HIV-2_287_ challenge (aqua line). Note, vaginal antibody is higher after immunizing exposures in experimental four animals compared to two controls evaluated, likely due to greater number of intravaginal SIV exposures (10 vs. 2) in these animals. **B**. Panels show HIV-2_287_ serum antibody levels (top), viral load by co-culture (second from top), HIV-2 RNA load (third from top; limit of detection above gray block), and CD4 cell concentrations (bottom) for control (blue lines) and experimental (red lines) macaques; tenofovir administration (pink block), and one HIV-2_287_ challenge (black line). Animals “immunized” with HIV-2_287_ demonstrated protection from disease (not infection) on challenge with homologous virus, with significantly lower viral infectivity (P=0.05) and no CD4 cell loss compared to controls (P=0.01).

Control animals not administered tenofovir prophylaxis all became infected by the intravaginal inoculations. Weekly blood sampling initially detected viral DNA by PCR at Study Week 4, 5, 2 and 2 in the four the SIV_mne_ control animals and at Study Week 2 in all four HIV-2_287_ control animals. Once infected, virus was consistently detected throughout the study in the control animals by co-culture and RNA and DNA PCR. CD4 cells decreased rapidly in the HIV-2_287_ infected control animals (**Figure 2B**).

### Induction of antibodies by exposure to virus during tenofovir prophylaxis

Exposure of the experimental animals to virus during tenofovir prophylaxis induced specific, non-neutralizing antibody in sera to SIV_mne_ or HIV-2_287_. The antibody concentrations in uninfected experimental animals were similar to the infected control animals (**Figure 2**). Vaginal antibodies to SIV were compared between the tenofovir-treated and control animals during study weeks 1-20, and the tenofovir-treated animals appeared to achieve higher levels (**Figure 2A**).

Neutralizing antibodies were not induced by the vaginal exposures during tenofovir prophylaxis assessed in sera at Study Week 20 just prior to viral challenge. However, neutralizing antibodies were detected in infected control animals (data not shown), and in all experimental macaques once infected by a challenge virus.

### Cellular immune responses after exposure to virus during tenofovir prophylaxis

SIV/HIV-specific cellular responses (IFN-γ ELISpot to Gag) were not induced in the experimental animals by viral exposures during tenofovir prophylaxis. ELISpot responses were detected in control animals following infection, and in experimental animals after infection with challenge viruses (data not shown).

### Viral challenge of SIV_mne_-immunized macaques

At Study Week 20, (six weeks after tenofovir was stopped, ten weeks after the final intravaginal inoculation during chemoprophylaxis), the SIV_mne_-immunized animals were challenged with 10^4^ TCID_50_ of SIV_mne_ (**Figure 2A**). Importantly, the SIV_mne_ used for intravaginal immunization and through the first challenge were grown in each animal’s autologous cells to avoid clearance based on allo-antigens incorporated into the viral envelope. Following this first challenge, one (M97032) of the four tenofovir-treated animals became infected. SIV DNA was persistently detected in the PBMC from this macaque beginning at Study Week 22 (two weeks post-intravaginal challenge), although viral co-cultures of this animal’s PBMC were rarely positive. After the initial challenge, SIV_mne_ was not detected in the other three macaques’ PBMC; neither by DNA PCR (6 µg DNA across 4 specimens/animal) nor by weekly co-cultures (6 cultures/animal).

The three uninfected experimental animals were challenged a second and third time with 10^4.5^ TCID_50_ SIV_mne_ grown in allogenic animals at Study Week 41 (21 weeks after the first challenge, 31 weeks after the last immunizing exposures during tenofovir prophylaxis). Two inoculations, 24 hours apart, with a greater quantity of virus, were administered due to the observation that the first two SIV_mne_ control macaques given weekly intravaginal inoculations of 10^4^ TCID_50_ did not have virus detected in their PBMC within the expected time frame. Virus is typically detected one to three weeks following intravaginal inoculation of macaques with 10^4^ TCID_50_ SIV_mne_, (unpublished, WaNPRC). One control animal had virus first detected by DNA PCR and culture five weeks following initial inoculation, the other had virus detected after four weeks by PCR and after five weeks by culture. This delay in the detection of virus suggested decreased infectivity of the inoculum. Several explanations were considered. First, while 10^4^ TCID_50_ was 10 times the 100% animal infectious dose in previous titration experiments, these data were limited to two animals per dose level (WaNPRC, unpublished), and infection by the intravaginal route is known to be relatively inconsistent [43]. Second, the virus could have been attenuated by the additional passage in the macaques’ own PBMC. Finally, the absence of disparate MHC antigens on the viral envelope could reduce infectivity as it is likely that fewer intravaginal lymphocytes became activated. Therefore, a greater amount of SIV_mne_ (two doses of 10^4.5^ TCID_50_ given 24 hours apart), and with virus grown in allogenic animals, was used for the second and third challenge of the three uninfected experimental animals. Two additional control animals were similarly challenged with the pool of virus grown in allogenic animals, with SIV_mne_ infection detected two weeks following the inoculation, these control animals had detectable viremia by both culture and DNA PCR.

After the second and third challenge in the experimental group, one (T97033) of the three macaques became infected (**Figure 2A**). SIV was detected in co-culture and by DNA PCR two weeks after the challenge and persistently thereafter. The two uninfected macaques each had seven negative PBMC cultures and DNA PCR (total of 7 µg PBMC/animal) from blood collected every other week over a period of 13 weeks. These two animals were subsequently challenged and infected with HIV-2_287_ (**Figure 2A**). The viral loads and CD4 cell counts differed between these two animals, one (K97255) had attenuated disease, with low levels of plasma HIV-2 RNA and normal CD4 counts.

### Viral challenge of HIV-2_287_-immunized macaques

At Study Week 20, ten weeks following the last immunizing HIV-2_287_ intravaginal inoculation (six weeks following the final dose of tenofovir), the experimental macaques were challenged with HIV-2_287_ grown in autologous PBMCs. All four immunized animals became infected. While HIV-2_287_ RNA loads appeared similar, disease was attenuated in three of the four animals compared to controls, with lower HIV-2_287_ loads in quantitative co-culture (P= 0.05), and delayed loss of CD4 cells (P= 0.01) (**Figure 2B**).

## Discussion

Our strategy to immunize macaques to a specific strain of virus by exposure to live SIV_mne_ or HIV-2_287_ during protective antiretroviral prophylaxis with tenofovir resulted in the generation of specific antibodies by all animals. Half of the SIV_mne_ immunized animals appeared to resist challenge with homologous virus. While the HIV-2_287_ immunized animals did not appear to resist infection, their disease was attenuated with lower concentrations of infectious virus and maintenance of normal CD4 cells counts. This macaque model suggests that partial protection to HIV could be achieved by exposure to infectious virus during antiretroviral prophylaxis.

The mechanism by which our immunization strategy may have blocked virus transmission to some animals and limited viral replication in others is uncertain. Tenofovir prophylaxis would have allowed virus to enter the animals’ cells but block reverse transcription of viral RNA necessary to establish infection. Virus-specific immune responses induced during tenofovir prophylaxis included binding antibodies in both the serum and vaginal secretions, but no detectable neutralizing antibodies or virus specific cellular responses.

We speculate that when the macaques were later challenged with live homologous virus, the binding antibodies may have contributed to protection in the SIV_mne_-exposed animals and to attenuation of disease in the HIV-2_287_ group. The virus specific IgG may have partially protected macaques from infection by antibody dependent cellular toxicity [44]. Vaginal secretory IgA has correlated with protection of high-risk humans from HIV infection [45, 46], and macaques [47], which supports our contention that vaginal IgA may have blocked infection by binding virus. However, serum IgA was shown to antagonize the effects of IgG-mediated ADCC in other studies [21, 48]; and vaginal secretory IgA has not been observed in all exposed-uninfected individuals [30, 49-51].

In the animals with attenuated disease, antibodies may have had diminished viral infectivity, as antibodies appear to modulate the level of viremia after the first month of infection [52]. Passive transfer of neutralizing IgG in 1-month-old HIV-infected rhesus macaques was reported to reduce plasma and PBMC-associated viremia [53]. Alternatively, allelic differentiation in viral antigen presentation due to genetic diversity, observed in rhesus macaques [54] (reviewed in [55]), may have contributed to attenuation of disease.

We did not assess whether the presence of viral proteins may have induced macrophages, dendritic cells or natural killer cells to secrete chemokines that block CCR5 or CXCR4 viral co-receptors or down-regulate the CCR5 receptor [56]. While high serum concentrations of RANTES [57] or vaginal application of CCR5 inhibitors has been demonstrated to block infectious intravaginal challenges [58, 59], if antiviral chemokines were induced in our animals, these would have had to persist for 10-20 weeks following the period of immunization to affect the outcome of the viral challenges.

While non-replicating virus presented by dendritic cells can induce CD4 and CD8 T-cell responses [60], it appears that there was insufficient processing and presentation of viral antigens to induce specific cellular immune responses in our experimental macaques. In a rhesus macaque model of intermittent PrEP with multiple low-dose rectal SHIV(SF162P3) challenges, virus-specific CD4 and CD8 T cells were detected in two of four animals protected from infection, but virus-specific antibodies were not found in these animals [61]. Live attenuated SIV vaccines have protected macaques against mucosal, but not intravenous, challenge, which suggests different responses might be needed to protect from mucosal compared to intravenous challenge [62]. While assays using PBMC did not reveal cellular immune responses, it is conceivable that dendritic cells in the vaginal mucosa initiated specific cellular immune responses limited to the mucosa or regional nodes [63]. The nodes draining the vagina of our experimental animals were not examined for viral nucleic acids, proteins or cellular responses, as these are difficult to access surgically. Also, it is possible that systemic cellular responses were diminished as has been noted to occur when mucosal immune responses are induced [64].

Two major challenges to vaccine development are the induction of immune responses towards viral strains that the host encounters and induction of immune responses that prevent infection. Live vaccines, such as live-attenuated intranasal influenza vaccine or virus-like-particles have induced greater mucosal IgA and cellular immune responses compared to intramuscular preparations and neutralize a more diverse repertoire of viruses and provide a survival advantage in animal experiments [65]. However, because live attenuated SIV/SHIV/HIV vaccine viruses cause persistent infection and can progress to severe immunodeficiency [66], the general approach has been to use vaccine vectors with a limited number of replication cycles to stimulate immune responses. Similarly, our strategy allows virus to enter the host cell, and while the reverse transcriptase inhibitor tenofovir prevents infection, vaginal IgA was induced.

One limitation of our study is the small sample size, which precluded highly statistically significant findings. In addition, non-human primate models using SIV and HIV-2 do not accurately mirror human infection with HIV. Atraumatic introduction of virus into the vagina, used in this model system, differs from sexual intercourse where trauma [67] or ulcerations may perturb mucosal barriers [68], which could overcome binding antibody. The animals we studied were healthy, while humans frequently have sexually transmitted infections that could increase lymphocytes that serve as viral targets in the mucosa surface [69].

The safety of our immunization strategy is uncertain. Pre-exposure prophylaxis has been noted to fail [70]. Vaginal immunization could increase the concentration of target cells in the vaginal mucosa, which has been speculated to increase susceptibility to HIV infection [69]. However, it is important to note that the production of systemic antibody and attenuated disease in infected macaques indicates that the mucosal exposures did not induce tolerance to viral antigens.

Similar to induction of antibody responses by intravaginal immunizations in the animals we studied, other studies have detected humoral and cellular immune responses after intravaginal [71, 72], rectal [73, 74] and nasal [75-78] administration of HIV/SIV. These observations suggest that administering prophylaxis during HIV exposures may effectively immunize individuals discordant for HIV infection, such as monogamous sexual partners and/or breastfeeding infants.

In summary, we explored a novel strategy to immunize macaques to lentiviruses. This model is relevant to sexual partners discordant for HIV infection and breastfeeding infants, who could benefit from protective immunity to homologous virus. Observational follow-up of humans, particularly breastfeeding infants completing pre-exposure prophylaxis and exposed to homologous viral strains could help evaluate findings of this pilot project. Clearly, additional studies are needed to define the mechanisms conferring partial protection to viral challenge, and to explore modalities to increase resistance to infectious virus.

## Acknowledgement

This project was supported by United States Public Health Service Grants, including R01 RR13273 (LMF), AI30034 (DA, MA, SH), and RR-00166 (WaNPRC).

We thank Gilead Sciences for providing tenofovir for this study.

